# Multimerization of Zika Virus-NS5 causes a ciliopathy and forces premature neurogenesis

**DOI:** 10.1101/719625

**Authors:** Murielle Saade, Diego S Ferrero, José Blanco-Ameijeiras, Elena Gonzalez-Gobartt, Victor M Ruiz-Arroyo, Elena Martínez-Sáez, Santiago Ramón y Cajal, Nuria Verdaguer, Elisa Martí

**Affiliations:** Developmental Biology Department, Instituto de Biología Molecular de Barcelona (IBMB-CSIC), Parc Científic de Barcelona, C/Baldiri i Reixac 20, Barcelona 08028, Spain; Structural Biology Department, Instituto de Biología Molecular de Barcelona (IBMB-CSIC), Parc Científic de Barcelona, C/Baldiri i Reixac 20, Barcelona 08028, Spain; Department of Pathology, Vall d’Hebron University Hospital, Translational Molecular Pathology, Vall d’Hebron Institute of Research (VHIR), Universitat Autònoma de Barcelona and Spanish Biomedical Research Network Centre in Oncology (CIBERONC)

**Keywords:** CNS growth, Neural Progenitor Cell, Zika Virus, Microcephaly, Centrosome, Ciliopathy

## Abstract

Zika virus (*ZikV*) is a flavivirus that infects neural tissues, causing congenital microcephaly. ZikV has evolved multiple mechanisms to restrict proliferation and enhance cell death, although the underlying cellular events involved remain unclear. Here we show that the ZikV-NS5 protein interacts with host proteins at the base of the primary cilia in neural progenitor cells, causing an atypical non-genetic ciliopathy and premature neuron delamination. Furthermore, in human microcephalic fetal brain tissue, ZikV-NS5 persists at the base of the motile cilia in ependymal cells, which also exhibit a severe ciliopathy. While the enzymatic activity of ZikV-NS5 appears to be dispensable, the Y25, K28 and K29 residues in the protein, that are involved in NS5-oligomerization, are essential for the localization and interaction with components of the cilium base, promoting ciliopathy and premature neurogenesis. These findings lay the foundation to develop therapies that target ZikV-NS5-multimerization, preventing the developmental malformations associated with congenital Zika syndrome

## Introduction

Zika virus (ZikV) is a flavivirus transmitted by the bite of the Aedes mosquito^1^. The 11 kb positive-sense, single-stranded RNA genome of ZikV encodes ten mature viral proteins: three structural proteins (C, prM and E) and seven non-structural proteins (NS1, NS2A, NS2B, NS3, NS4A, NS4B, and NS5)^1^. A single amino acid substitution (S139N) in prM seems to be responsible for the change in virus tropism that provokes a dramatic increase in neonatal microcephaly^22^, and that has led to this virus being declared a global threat to public health. ZikV can directly infect human neural progenitor cells (NPCs) in both 2D and 3D *in vitro* models of the developing cerebral cortex, resulting in defects resembling congenital microcephaly^3–8^. However, the mechanisms by which ZikV disrupts neurogenesis have not yet been fully elucidated. Moreover, macaque monkey infection provokes regional disturbances in the brain that impair postnatal neurogenesis, provoking cognitive deficits and epilepsy^9^. This vulnerability of late neurogenic regions to ZikV also persists in adult mice^10^. As it is well known that human brain development extends for many years beyond birth and that adult neurogenesis influences cognitive functions, it is no longer safe to consider ZikV as a transient infection in adult humans without marked long-term effects.

The limited availability of human tissue to perform histological analyses at different developmental stages^11^ emphasizes the need to use *in vivo* animal models to understand congenital Zika syndrome. Here we used the chick embryo neural tube (NT) to screen non-structural (NS) ZikV protein components and assess their impact on neurogenesis, particularly given that this *in vivo* model has served to identify fundamental processes in mammalian neural development and human disease. Through this approach, ZikV-NS5 was seen to disturb the growth of the nervous system. ZikV-NS5 is the largest NS protein (~100 kDa) and it is essential to the viral life cycle. It contains two functional domains, a N-terminal methyltransferase (MTase) domain responsible for the catalysis of 5’-mRNA capping and methylation, and a RNA-dependent RNA polymerase (RdRP) domain at the C-terminus that drives genome replication^12–15^. ZikV-NS5 can form dimers and higher-order oligomers that regulate its enzymatic functions, as well as its association with other viral or host factors during infection, making it an extremely attractive anti-viral target^13^.

To understand how ZikV-NS5 affects neurogenesis, we searched for the host ZikV-NS5 interactome in human NPCs in a yeast two-hybrid (Y2H) assay. This unbiased approach detected previously identified ZikV-NS host interactors^16,17^ and it revealed that the ZikV-NS5 protein interacts with cellular components known to localize at the base of the cilium. We showed that primary cilia elongation is impaired in NPCs expressing ZikV-NS5 and that neural delamination is accelerated, two key events that promote premature neurogenesis and impair CNS growth. However, these effects were hindered when ZikV-NS5 multimerization was prevented by point mutations. In addition, we observed a severe ciliopathy in the ependymal cells (ECs) that line the ventricular cavities of a ZikV-infected human fetal microcephalic brain, a phenomenon that might impair the cerebrospinal fluid (CSF) flow necessary to maintain brain homoeostasis and for toxin washout. Together our data indicate that the ZikV-NS5 causes ciliopathies – both a motile ciliopathy and a primary ciliopathy- which contribute to the collection of developmental malformations provoked in congenital Zika syndrome, such as microcephaly and ventriculomegaly.

## Results

### ZikV-NS5 interacts with basal ciliary proteins in NPCs

To investigate whether individual non-structural proteins (NS1-5) encoded by ZikV can directly regulate neurogenesis (Fig. 1a), we monitored the expression of the anti-proliferative gene Tis21 (PC3, BTG2)^18^ 24 hours after electroporation (hpe) of constructs encoding each of these proteins into Hamburger and Hamilton (HH) stage 12 chick embryo NTs^19^. Each ZikV-NS protein was co-electroporated with the pTis21-Luciferase reporter gene and the expression of all the ZiKV proteins in early NPCs was monitored (Extended Data Figs. 1a-e). ZikV-NS2 and ZikV-NS5 provoked an increase in luciferase activity (Fig. 1b) and indeed, ZikV-NS2 has been reported to dampen NPC proliferation by directly interacting with components of the adherens junction (AJ) complex^20^. The expression of ZikV-NS5 also reduced the expression of the progenitor marker pSox2, enhancing the expression of additional pan-neural markers like pNeuroD and pTubb3 (Fig. 1c, Extended Data Figs. 1e-f), without disrupting NT tissue integrity or affecting cell viability (Extended Data Figs. 1g-j).

**Figure 1.**
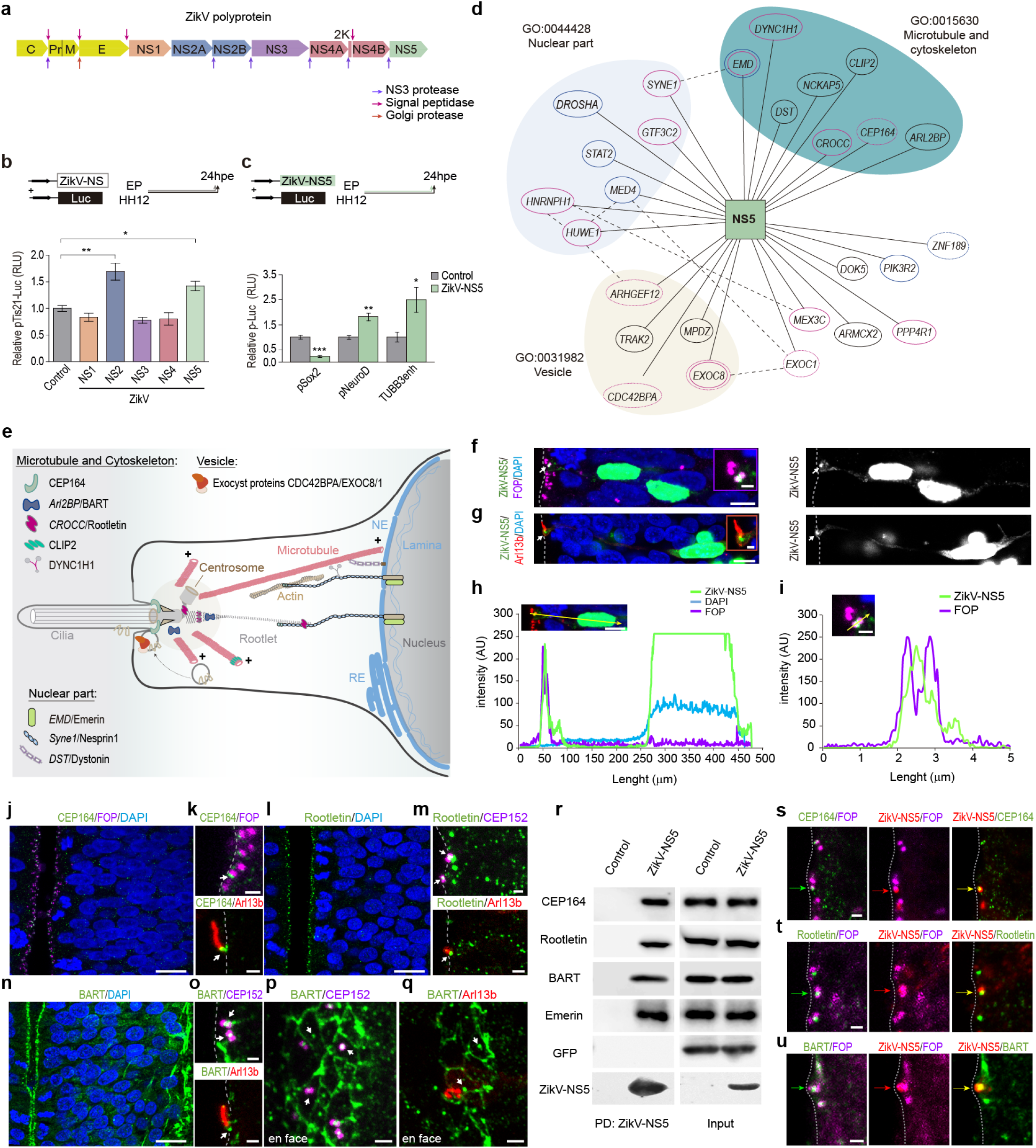
ZikV-NS5 interacts with proteins at the base of cilia. **(a)** The scheme depicts the structural (yellow) and non-structural (NS) proteins of the Zika virus polyprotein, the arrows indicate the sites of proteolytic cleavage. **(b)** Quantification of the Luc/Renilla activity of the pTis21-Luc reporter after electroporation of the ZikV-NS DNAs indicated (plots show the mean ± s.e.m., n=6-8 embryos/condition) **(c)** Quantification of the Luc/Renilla activity of the pSox2-Luc, the pNeuroD-Luc and the Tubb3enh-Luc reporters after electroporation of ZikV-NS5 (plots show the mean ± s.e.m., n=6-8 embryos/condition) **(d)** Network representation of the ZikV-NS5-host interactome in human NPCs, where the bait protein (NS5) is represented as a square and the 27 interacting host proteins as ellipses. The subcellular localization (colour of nodes) of the host proteins are taken from Gene Ontology. Proteins previously reported to interact with ZikV-NS5 in neural cells are in pink and those that interact in HEK291 cells are in blue. Published physical interactions of the host proteins are indicated by dotted lines. **(e)** The scheme depicts an overview of the apical pole of a NPC where the primary cilium extends into the NT lumen. The centrosome at the cilium base organizes microtubules that link the nuclear envelope (NE) to the cilium base through the Rootlet. Key to proteins that appeared to interact with ZikV-NS5 in the Y2H screen. **(f,g)** Selected images show the subcellular distribution of ZikV-NS5 in electroporated NPCs. **(f)** ZikV-NS5-FLAG visualised by anti-FLAG staining (green) in the nucleus (DAPI, blue) and in the centrosomes (arrow) lining the NT lumen labelled with anti-FOP (purple). **(g)** ZikV-NS5-FLAG located at the base of the cilium (arrow) labelled with Arl13b-RFP (red). **(h,i)** Plots of the fluorescence intensity (AU) at the distances from the NT lumen (zero): purple labelled centrosomes, blue labelled nuclei and green labelled ZikV-NS. **(j,k)** Representative image of a NT section showing endogenous CEP164 (green) lining the NT lumen and arrows pointing to the distal co-localization in the mother centriole with FOP (purple) at the base of the Arl13b-RFP labelled cilium (red). **(l,m)** Section showing endogenous Rootletin (green) in which the arrows indicate its co-localization with the centrosome marker CEP152-GFP (purple) at the base of the Arl13b-RFP labelled cilia (red). **(n,o)** Arrows point to endogenous BART (green) co-localizing with CEP152-GFP (purple) at the base of the Arl13b-RFP labelled cilia (red). **(p,q)** *En face* imaging of the BART labelled apical belt of neuroepithelial end-feet, with arrows pointing to the co-localization with CEP152 labelled centrosome (purple) and at the base of Arl13b-RFP labelled cilia (red). **(r)** HEK-293 cells were co-transfected with ZikV-NS5-GFP and the DNAs indicated, and the cell extracts were analysed by western blots. Pull-down (PD) assays show ZikV-NS5 binding to Cep164, Rootletin, BART and Emerin, with GFP serving as a negative control. **(s-u)** Representative images of ZikV-NS5-FLAG (red) co-localization with the FOP labelled centrosome (purple) and with endogenous CEP164 (green, **s**), endogenous Rootletin (green, **t**) and endogenous BART (green, **u**)lining the NT lumen (dotted line). The proteins are indicated with coloured arrows according to their fluorescence labelling. .\*\*\**p*<0.001, \*\**p*<0.01; exact *P* values 0.0091 (**b**), 0.0079 (**c**), \**P*<0.05 exact *P* values, 0.0416 (**b**), 0.0101 (**c**), ns: not significant. **b**, one-way ANOVA; **c**, two-sided unpaired t-test. Scale bars 5 μm (**f, g** inset 1μm), 9 μm (**j, l, n**), 1 μm (inset **f**, inset **g**, **k, m, o, s-u**), 2 μm (**p**, **q**).

We searched for the cellular proteins that might interact with ZikV-NS5 and influence neurogenesis. As such, the ZikV-NS5 (1-904)-LexA fusion protein was used as bait in a Y2H screen of a human foetal brain library (HFBR_RP1_hgx4776v1_pB29). After analysing 99.6 million interactions and processing more than 141 colonies, a total of 27 ZikV-NS5-host interactors were defined (Supplementary Table 1), ~70% of which were common to those interactors identified previously^16,17^ (Supplementary Table 2). Among the host proteins detected, enrichment of the microtubule/cytoskeleton, vesicle and nuclear compartments was revealed by gene ontology (GO) analysis (Figure 1d). Interestingly, several of these proteins are known to localize to the base of the cilium and/or, to link the nuclear envelope to the cytoskeleton (Fig. 1e; Supplementary Table 1).

A hallmark of NPCs is the single primary cilium at their apical surface, which is nucleated by the basal body where the mother centriole is docked to the plasma membrane through its appendage proteins^21^ (Fig. 1e). This cilium extends into the NT lumen, where it is able to detect the growth factors in the CSF that control CNS growth^22^. Findings from the interactome prompted us to study the subcellular distribution of ZikV-NS5 in NPCs by introducing a plasmid encoding ZikV-NS5-FLAG tagged into the chick embryo NT. As expected given the nuclear localization signals encoded in this protein, the NS5-FLAG protein accumulated in the nucleus^12^ (Figs. 1f-h), a subcellular distribution unique among the ZikV proteins (Extended data Figs. 1a-e). However, some ZikV-NS5 also accumulated at the base of the cilia, which was identified through the small GTPase ADP-ribosylation factor-like 13b fused to the red fluorescent protein (Arl13b-RFP) that specifically associates with the ciliary membrane^23,24^. ZikV-NS5 co-localizes with the centrosome, identified as a pair of dots at the surface of the ventricular zone (VZ) where the centrosome proteins 152K (CEP152)^25^ and FGFR1 Oncogene Partner (FOP) ^26^ were detected (Figs. 1f-i). Moreover, the docking of ZikV-NS5 to the centrosomes persisted throughout mitosis in these NPCs (Extended Data Fig.1k-n).

To confirm that ZikV-NS5 interacts directly with components of the cilium base in a cell context, endogenous centrosome protein 164K (CEP164) was detected in NPCs by immunofluorescence, lining the NT lumen where it co-localizes with the FOP-labelled mother centriole at the cilium base (Figs. 1j,k). CEP164 is a mature centriole-specific protein that localizes to the basal body distal appendages and it is required for assembly of the primary cilium^27^. ZikV-NS5 pulled down CEP164, in HEK293 cell extracts (Fig. 1r, and Extended Data Figs. 1r,) and in NPCs, ZikV-NS5 co-localized with CEP164 and with centriole markers (Fig. 1s, Extended Data Fig. 2d). In NPCs, the endogenous rootlet coiled-coil protein (*CROCC*/Rootletin) was detected by immunofluorescence in the rootlet fibres, as well as lining the NT lumen where it co-localized with centrosomal markers at the cilium base (Figs. 1l,m). Rootletin localizes to the intercentriolar linker and it is required for the correct positioning of the cilium basal body relative to the cell nucleus^28,29^. ZikV-NS5 also pulled down Rootletin in HEK293 cell extracts (Fig. 1r, and Extended Data Fig. 1r,), and ZikV-NS5 co-localized with Rootletin and with centrosomal markers in NPCs (Fig. 1t, Extended Data Fig. 2e). Moreover, immunofluorescence demonstrated the presence of the endogenous ADP ribosylation factor-like 2-binding protein (*ARL2BP*/BART) in NPCs that line the NT lumen, where it co-localized with the centrosomal marker CEP152 at the cilium base (Figs. 1n-q). BART localizes to the basal body and it is required for cilia elongation^30^. *En face* NT imaging revealed BART also localized to the apical belt (Figs. 1p, q) and the base of the primary cilium identified by Arl13b (Fig. 1q). ZikV-NS5 pulled down BART in transfected HEK293 cells (Figure 1r, and Extended Data Figs. 1r) and in NPCs, ZikV-NS5 co-localized with BART and with centrosomal markers (Fig. 1u, Extended Data Fig. 2h). In addition, the Y2H screen retrieved three ZikV-NS5 protein interactors that localize to the nuclear envelope (Emerin, Nesprin and Dystonin: Fig. 1e). ZikV-NS5 pulled down Emerin from HEK293 cells (Fig. 1r), a member of the nuclear lamina-associated protein family that controls anchorage to the cytoskeleton and that links centrosomes to the nuclear envelope through an association with microtubules ^31,32^. Together these data indicate the ZikV-NS5 interacts with protein components of the base of the primary cilium in NPCs.

### ZikV-NS5 causes a ciliopathy and premature neuron delamination

In dividing NPCs, cilia length shortens prior to entry into mitosis and the ciliary membrane remnant is internalized along with the mother centrosome. The daughter cells that inherit the mother centrosome can reform long cilia faster and they remain as NPCs, while the differentiating daughter cells delaminate from the ventricular surface and migrate to the lateral NT (Fig. 2a)^23,33^. We assessed whether ZikV-NS5 affects cilia length, measuring the length of Arl13b-RFP labelled cilia protruding from the two FOP-labelled centrioles at the VZ surface in NPCs. In control cells the cilia length varied as a function of the cell cycle phase and nuclear position (median=2.08 ± 1.27 μm, n=100), while the cilia were significantly shorter in ZikV-NS5 electroporated NPCs (median=0.69 ± 0.3 μm, n=55: Figs. 2b-d; Extended Data Figs. 2a-c). Co-expressing all ZikV associated NS proteins in NPCs did not affect the localization of NS5 to the cilia base or revert the ciliopathy (Extended Data Figs. 1o-q). To search for the mechanism by which ZikV-NS5 mediates cilia shortening, we quantified the endogenous target proteins in NPCs. Compared to the controls, there was a significant reduction in CEP164 protein in ZikV-NS5 electroporated NPCs (control median=8.88 ± 3.29 AU, n=58; ZikV-NS5 median=1.68 ± 1.27, n=31: Figs. 2e, h; Extended Data Figs. S2d, f). Interestingly, less CEP164 was reported previously in ZikV-infected NPCs forming brain organoids^6^. In addition, there was less endogenous centrosomal Rootletin and BART in NPCs upon exposure to ZikV-NS5 (Rootletin control median=20.3 ± 11.12 AU, n=75; ZikV-NS5 median =5.1 ± 8.67 AU, n=37; BART control median=3.6 ± 4.7AU, n=114; ZikV-NS5 median=1.13 ± 4.4 AU, n=36: Figs. 2f,g, i, j; Extended Data Figs. 2e, g). Hence, the reduction in the endogenous CEP164, Rootletin and BART protein at the centrosome of cells expressing ZikV-NS5 correlated with the impaired cilia elongation in NPCs. In turn, cilia shortening might impair the reception of extracellular growth signals, such as that provided by Sonic Hedgehog (Shh), an important regulator of proliferation^33,34^.

**Figure 2.**
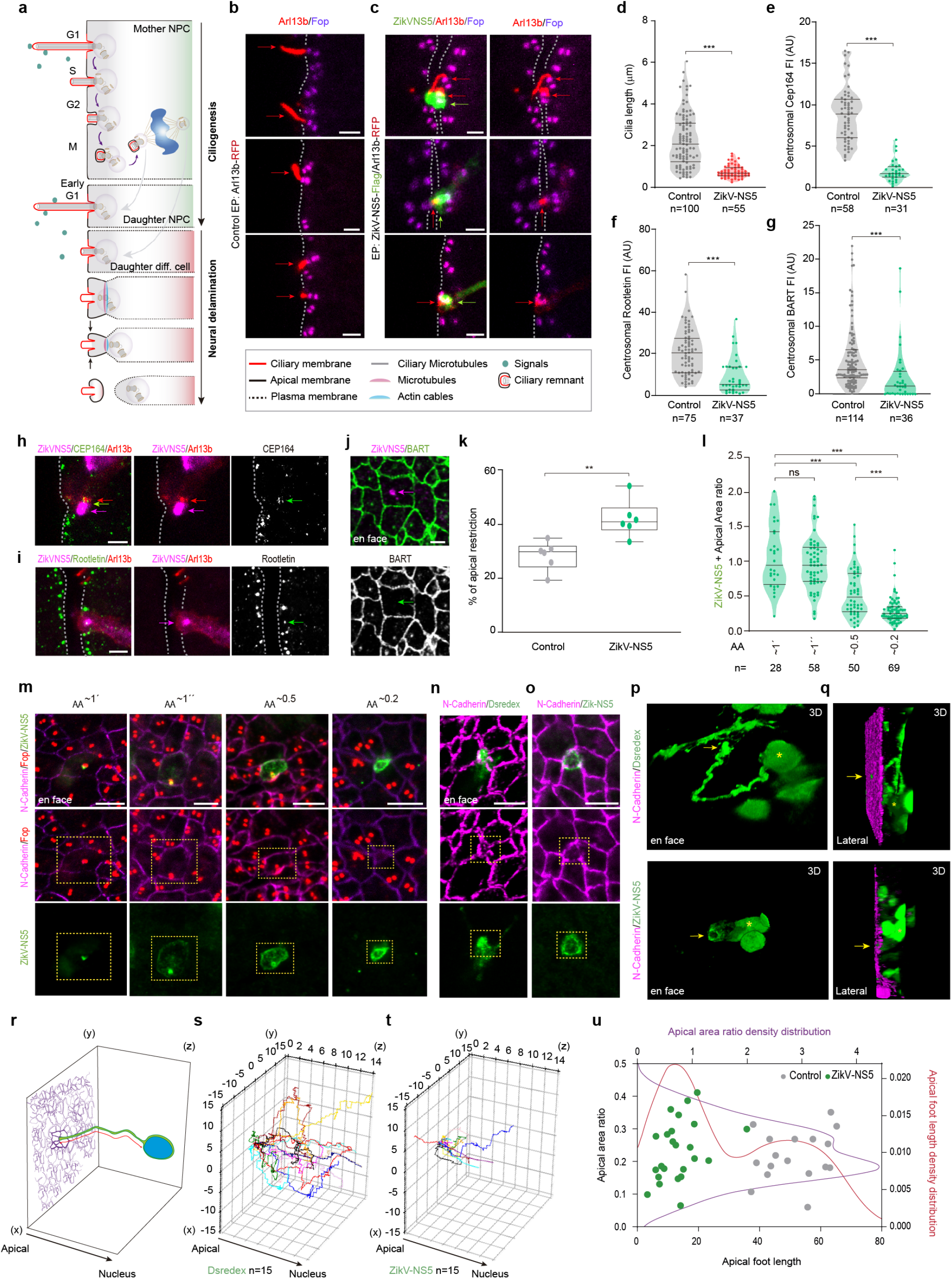
ZikV-NS5 localization to the cilium base impairs cilia elongation and promotes neural delamination. **(a)** The scheme depicts the variation in cilia length during the cell cycle in a dividing NPC. The daughter cell remaining as a NPC regrows an apical primary cilium, while the daughter cell entering differentiation delaminates from the epithelium. **(b, c)** Selected images show the Arl13b-RFP labelled cilia (red arrows) and FOP stained centrosomes (purple) lining the NT lumen (dotted line) in (b) control and (c) ZikV-NS5 electroporated (green arrows) NPCs. **(d)** Violin plots of cilia length in control and ZikV-NS5 electroporated NPCs **(e-g)** Violin plots of the fluorescence intensity (AU) of the indicated endogenous proteins at the centrosome in control and ZikV-NS5 electroporated NPCs. **(h,i)** Representative images of ZikV-NS5-FLAG (purple arrow) co-localization with Arl13b (red arrows) and endogenous CEP164 (**h**) or endogenous Rootletin (**i**) (green arrows) **(j)** *En face* images of endogenous BART labelled apical ring (green) and its absence from the centrosome where ZikV-NS5-FLAG is located (purple arrow) **(k)** Plots of the proportion of cells with apically restricted areas in control and ZikV-NS5 transfected NTs (n= 6 embryos; error bars correspond to the median ± S.D.) **(l)** Violin plots of the apical areas (AA) in relation to ZikV-NS5 positioning. The non-restricted areas (~1) correspond to ZikV-NS5 co-localization at the centrosome and the restricted areas (≤0.5) to ZikV-NS5 localization at the apical belt. **(m)** *En face* images of N-cadherin (purple) labelled AJs, FOP (red) labelled centrosomes and ZikV-NS5-FLAG subcellular localization (green) according to the apical area (AA). **(n,o)** *En face* images of N-cadherin (purple) showing restricted apical areas in Dsredex-control (**n**) and ZikV-NS5-FLAG electroporated NPCs. **(p,q)** 3D reconstructions of the apical foot in Dsredex-control or ZikV-NS5-FLAG transfected NPCs (green). Arrows point to the apical end foot area facing the NT lumen (N-cadherin, purple: asterisks correspond to nuclei). **(r)** The scheme depicts the apical foot length (red line) as the distance between the restricted apical-en foot area facing the NT lumen (purple) and the basal nucleus (blue) in NPCs. **(s,t)** 3D Python Plots of apical foot length in control (**s**) and ZikV-NS5 (**t**) transfected NPCs. **(u)** Plots of the apical foot length relative to the apical-end foot area restriction in control (grey dots) and ZikV-NS5 expressing NPCs (green dots). The density distribution of the apical-end foot area restriction presents a Gaussian distribution (in purple), while the density distribution of the apical foot length presents a bimodal distribution (Red). In violin plots, the upper and lower lines indicate the interquartile range, and the middle line the median. .\*\*\**p*<0.001, \*\**p*<0.01; exact *P* value 0.0043 (**k**), \**P*<0.05, ns: not significant. **d** -**k**, Mann-Whitney *U* test; **l**, Kruskal-Wallis test. Scale bars 2 μm (**b**, **c**, **h**-**j**) 4 μm (**m**-**o**)

In NPCs, apical centrosomes organize a microtubule ring that, when aligned with the actin cables, plays an active role in reducing their apical surface and withdrawing their apical endfoot from the ventricular surface (Fig. 2a)^35,36^. We evaluated whether ZikV-NS5 might influence neural delamination, since BART localizes to this apical belt (Figs. 1p-q, 2j; Extended Data Fig. 2h, j-k). We measured the apical area (AA) of electroporated cells surrounded by non-electroporated neighbors in *en face* images of the NT, the latter used as a reference for AA restriction. The proportion of NPCs with a smaller AA is higher following ZikV-NS5 electroporation (control ratio=28.32 ± 5.13; ZikV-NS5 ratio=41.90 ± 6.7: Fig. 2k; control median=0.7 ± 0.53 AU, n=163; ZikV-NS5 median =0.51 ± 0.46 AU, n=213: Extended Data Fig. 2l). The AA was delimitated by the AJ-associated protein N-cadherin and the centrosome was defined by FOP (Fig. 2m). In the non-restricted AA, ZikV-NS5 localizes to either the central or lateral positioned centrosome, a feature that precedes neural delamination (AA~1’ ratio=0.92 ± 0.46; AA~1” ratio=0.92 ± 0.39: Figs. 2l-m, Extended Data Fig. 2i ^37^). However, the localization of ZikV-NS5 to the apical belt was correlated with a reduction in the AA (AA~0.5 ratio=0.46 ± 0.35; AA~0.2 ratio=0.23 ± 0.19: Figs. 2l-m, Extended Data Fig. 2l), coinciding with early apical to basal migration of the centrosome (Figure 2m, Extended Data Figs. 2i, m-n, Video 1). Moreover, a reduction in the AA at the ventricular surface is preceded by the basal migration of the new-born delaminating neuronal cell body^35^. Confocal images of Dsredex or ZikV-NS5 electroporated cells were used to generate 3D reconstructions and to track the cell processes that extend from the apically endfoot restricted surface to the nucleus (Figs. 2n-r). ZikV-NS5 electroporated cells have a short apical foot length (control length=51.5 ± 9.85μm, n=17; ZikV-NS5 length=13 ± 7μm, n=25: Figs. 2 o-q, t, u, Videos 2-4) compared to control (Figs. 2n, p, q, s, u, Videos 5, 6), indicating that ZikV-NS5 promotes apical endfoot restriction at the ventricular surface before the basal migration of the new-born delaminating neuronal cell body is completed.

Finally, when we assessed the expression of Tis21 to determine if these cellular events promote terminal neurogenesis, reporter activation was associated with the localization of ZikV-NS5 to the cilium base (Extended Data Fig. 2o). Quantification of Tis21p-RFP^+^ NPCs at 16 hpe (n=5 embryos) showed that 93 ± 7% of the NPCs expressed pTis21-RFP when ZikV-NS5 was localized to the cilium base, as opposed to 25 ± 10% of NPCs in which ZikV-NS5 localizes to the nucleus (n=5 embryos), or 25 ± 3% of the control NPCs (n=6 embryos: Fig. 2p). Together, these data indicate that primary cilia elongation is impaired and apical enfoot restriction is accelerated in NPCs exposed to ZikV-NS5 protein, two key events that promote neural delamination. These results prompted us to study how the ZikV-NS5 protein affects primary neurogenesis and embryonic CNS growth.

### The multimeric arrangement of ZikV-NS5 is required to promote terminal neurogenic divisions

Normal growth of the CNS requires a fine balance between the different modes of NPC division and a premature switch to neuron generating divisions may cause microcephaly^33^. Symmetric proliferative divisions that leave two NPC daughter cells (PP) can be monitored *in vivo* with the pSox2-EGFP construct, whereas pSox2-EGFP and the neurogenic pTis21-RFP reporter are co-expressed in asymmetric divisions that produce one NPC and one differentiating neuron (PN). By contrast, symmetric neurogenic divisions generating two neurons (NN) that detach from the ventricular zone can be monitored by the expression of the pTis21-RFP reporter (Fig. 3a)^24,38,39^. To determine whether ZikV-NS5 affects the mode of NPC division, stage HH10/12 embryos were co-electroporated with ZikV-NS5 together with the pSox2-EGFP/pTis21-RFP reporters (Fig. 3b). When the embryos were analyzed at 24 hpe through fluorescent activated cells sorting (FACs), the rate of PP divisions (pSox2-EGFP^+^ cells) was seen to decrease significantly in ZikV-NS5 embryos (from 51 ± 2% in controls to 30 ± 2.8% in ZikV-NS5 embryos). This reduction in PP divisions took place at the expense of an increase in neurogenic divisions (from 18 ± 1.6% in controls to 46.4 ± 5.6% in ZikV-NS5 embryos: Fig. 3c). Similarly, when embryos electroporated were immunostained at 24 hpe, there was again a clear decrease in the rate of PP divisions (in pH3^+^ cells) in ZikV-NS5 embryos (from 32± 3.4% in the controls to 16± 6.5% in ZikV-NS5-EP embryos), which was compensated by an increase in NN divisions (from 13± 3.4% in controls to 33 ± 6.4% in ZikV-NS5-EP embryos: Figs. 3d-f). Together these data suggest that by preventing cilia elongation, the ZikV encoded NS5 protein alone is sufficient to switch the mode of NPC division and to promote neural delamination. Thus, ZikV-NS5 drives the exhaustion of the neural progenitor pool.

**Figure 3.**
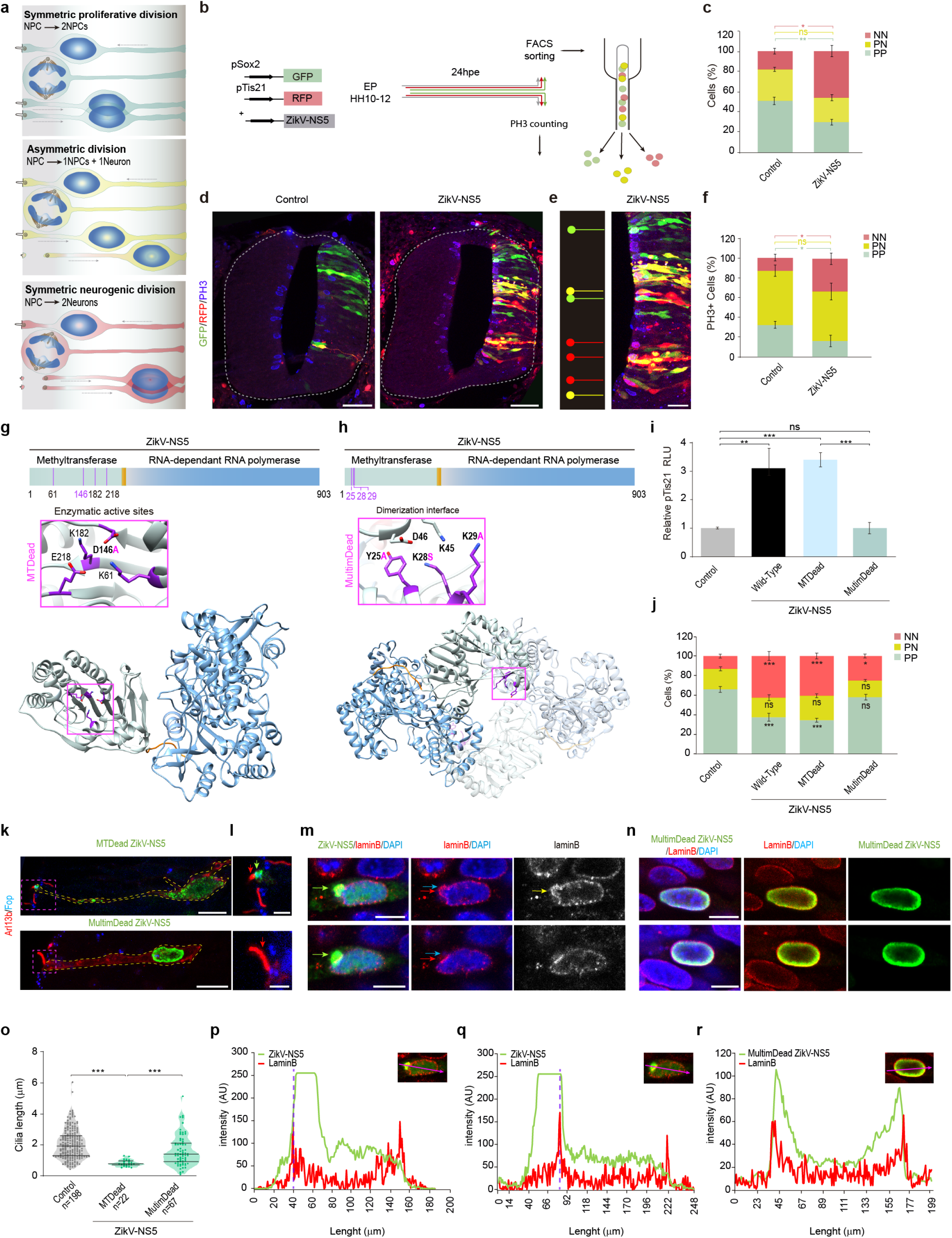
The multimeric arrangement of ZikV-NS5 is required for its localization to the cilia base and the promotion of terminal neurogenic divisions. **(a)** Scheme showing the three modes of NPC division. **(b)** Scheme representing the reporter co-electroporation experiments, harvested at 24 hpe, either for flow cytometry analysis (FACS) or for anti-PH3 immunostaining. **(c)** Quantification of the cells expressing the reporter in each condition: PP (pSox2^+^/pTis21^−^, green), PN (pSox2^+^/pTis21, yellow), and NN (pSox2^−^/pTis21^+^, red). The data represents the mean ± s.e.m). **(d,e)** Selected images show pSox2 (green) and pTis21 (red) NPCs, and pH3 stained mitoses (blue) in control or ZikV-NS5 (e) electroporated chick embryo NTs. Green, yellow and red lines in **e** indicate mitotic PP, PN and NN divisions, respectively. **(f)** Quantification of reporter-expressing pH3^+^ dividing cells in each condition. The data represents mean ± s.e.m.: \*\*\**P*<*0.001* \*\**P*<*0.0*, \**P*<*0.05*. **(g)** The scheme depicts the MTase (light blue) and RdRp (dark blue) domains in the ZikV-NS5 protein. 3D representation of one NS5 monomer shown in an orientation from the top of RdRP with the MTase domain shown in grey and the RdRP finger, palm and thumb sub-domains in blue. Inset showing the amino acids involved in the enzymatic activity, with the amino acid substitution highlighted in purple. **(h)** The scheme depicts the ZikV-NS5 protein and a 3D representation of one zikV-NS5 dimer. Inset shows the amino acids involved in dimerization, highlighting the amino acid substitution in purple. **(i)** Quantification of the Luc/Renilla activity of the pTis21-Luc reporter after electroporation of the DNAs indicated (plots show the mean ± s.e.m., n=6-8 embryos/condition). **(j)** Quantification of reporter-expressing FACS sorted cells in each condition: PP (pSox2^+^/pTis21^−^, green), PN (pSox2^+^/pTis21^+^, yellow) and NN (pSox2^−^/pTis21^+^, red). The data represents the means ± s.e.m). **(k,l)** Representative images of Arl13b (red) at cilia, the FOP (blue) labelled centrosomes at the cilia base and the localization of the ZikV-NS5 variants: ZikV-NS5-MTDead at the cilium base (green arrow in **l**), ZikV-NS5-MultimDead excluded from the cilium base (red arrows point cilia in **l**). **(m,n)** Representative images of nuclear ZikV-NS5_FLAG (green) in two separate z stacks. Upper image shows nuclear aggregates (green arrow) causing disruption of the nuclear envelope. Lower ‒image shows ZikV-NS5FLAG aggregates (green arrow) exit from the nuclear envelop. Nuclear Envelop disruption is highlighted by Lamin B (red arrow) and DAPI (blue arrow). ZikV-NS5-MultimDead shown in two separate cells in **n**, does not form aggregates and localizes at the vicinity of the nuclear envelop. **(o)** Violin plots of the cilia length in control NPCs and those electroporated with the mutant ZikV-NS5 constructs (the upper and lower lines indicate the interquartile range, and the middle line the median). **(p,q)** Plots of the fluorescence intensity (AU) of green labelled ZikV-NS5 relative to the red lamin B labelling at the two separate Zstacks shown in **m**. The purple dotted line depicts nuclear envelope disruption. **(r)** Plots of the fluorescence intensity (AU) of green labelled ZikV-NS5-MultimDead with the same intensity profile as the red labelled lamin B. .\*\*\**p*<0.001, \*\**p*<0.01; exact *P* values 0.0054 (**c**) 0.0034 (**i**), \**P*<0.05; exact *P* values 0.0129 (**c**) 0.0151 (NN, **f**) 0.0398 (PP, **f**), 0.0225 (**j**).ns: not significant. **c**, **f** two-sided unpaired t-test; **i, j**, one-way ANOVA; **o**, Kruskal-Wallis test. Scale bars 30 μm (**d**), 15 μm (**e**). 7 μm (**k**), 2 =m (**l**), 5 μm (**m**, **n**).

To better define the neurogenic effects of ZikV-NS5, we monitored the expression of Tis21 after co-electroporation of the reporter and the ZikV-NS5 N-terminal (NTerm, residues 1–262; MTase) or ZikV-NS5 C-terminal (CTerm, residues 267–903; RdRP) domains into the chick embryo NT (Extended Data Figs.3a, b). In contrast to the ZikV-NS5-CTerm domain, the ZikV-NS5-NTerm alone was sufficient to enhance pTis21-Luciferase reporter activity (Extended Data Fig. 3b). Moreover, the ZikV-NS5-NTerm drove an increase in the rate of NN divisions in the embryos, as witnessed in immunostained embryos (pH3^+^ cells; Extended Data Fig. c-e) and by FACs analysis (Extended Data Figs. c, f) Together these data suggest that the neurogenic capacity of the ZikV-NS5 protein resides in its N-terminal domain, which prompted us to assess whether enzymatic activity was required to promote neurogenesis.

The MTase domain of the NS5 flavivirus protein mediates the addition of a guanine cap to the 5’ end of the RNA genome, and the sequential methylation of guanine N-7 and ribose 2’-O to form a type 1 cap structure (^m7N^GpppA^2’Om^). This activity is essential to evade the host’s immune response and for replication, increasing the viral polyprotein translation efficiency. The ZikV MTase active site contains residues K61, D146, K182 and E218, which are in close contact with the S-adenosylmethionine (SAM) co-factor, the methyl group donor for the reaction (Fig. 3g). However, a single amino acid substitution (D146A) was sufficient to abolish MTase activity^13^ (MTDead). To assess whether this enzymatic activity is required for the neurogenic capacity of ZikV-NS5, we co-electroporated the pTis21-Luciferase reporter together with the ZikV-NS5-MTDead construct. Consequently, the luciferase activity of the reporter increased in response to mutant ZikV-NS5 MTase in a similar manner to that produced by the ZikV-NS5 protein (Fig. 3i). Indeed, the ZikV-NS5-MTDead construct also provoked an increase in the rate of NN divisions in embryos when analyzed by FACs (13 ± 3% in controls, 42.5 ± 4.5% in ZikV-NS5, and 40.5 ± 3% in ZikV-NS5-D146A embryos: Fig. 3j).

We recently characterized the supramolecular arrangement of the full-length ZikV-NS5, highlighting the assembly of NS5 monomers into dimeric structures, as well as the interactions of dimers to form a higher order fibrillar structures^12^. The main interface for dimer formation involves the MTase-MTase contacts, connecting amino acids Y25, K28 and K29 in one ZikV-NS5 molecule with residues K45 and D46 of the second ZikV-NS5 subunit^12^ (Fig. 3h). To test whether the multimeric arrangement of ZikV-NS5 is required to promote premature neurogenesis, we used the ZikV-NS5-Y25A/K28S/K29A mutant that fails to form dimers^13^ (MultimDead). This MultimDead mutant protein did not increase pTis21-Luc reporter activity (Fig. 3i) and in embryos analyzed by FACs at 24 hpe, dimerization was required for ZikV-NS5 to promote the neurogenic mode of NPC division (25 ± 2% NN divisions in ZikV-NS5-MultimDead embryos: Fig. 3j). These data indicate that while the enzymatic activity of NS5 is dispensable, ZIKV-NS5 oligomerization is required for this protein to promote neurogenesis.

To define the cellular mechanisms altered by the Zikv-NS5 protein variants, we followed their subcellular distribution in chick embryo NPCs. The ZikV-NS5 MTDead protein was detected in the nucleus and at the cilium base, like the wild type protein, while the MultimDead protein variant was retained within the nucleus, concentrating in the vicinity of the nuclear envelope (Figs. 3k-n). Moreover, the multimerization of ZikV-NS5 contributes to the formation of ring-like nuclear aggregates that exit the nucleus toward the apical foot and localize at the centrosome forming the cilia base (Extended Data Fig. 3g). Furthermore, the nuclear exit of multimerized ZikV-NS5 is associated to the transient disruption of the nuclear envelope evident as Lamin B blebs and protruding chromatin (Figs. 3m, n, p-r). Indeed, this transient loss of nuclear integrity has already been reported in ZikV infection^40^, as well as in other viral infections^41^. Nuclear retention of the ZikV-NS5 MultimDead protein, might rely on its reduced capacity to bind the nuclear envelope components such as Emerin (Extended Data Figs. 3j,k). Hence, the trafficking of ZikV-NS5 to the cilia base appears to depend on its capacity to form multimers.

The retention of ZikV-NS5 within the nucleus was sufficient to prevent the ciliopathy, as evident by the length of the Arl13b-RPF labelled cilia in NPCs (control median=1.98 ± 0.97 μm, n=198; ZikV-NS5-MTDead median= 0.82 ± 0.2 μm, n=22; ZikV-NS5 MultimDead median= 1.44 ± 1 μm, n=67: Fig. 3o). Even though the ZikV-NS5 MultimDead protein variant has a reduced capacity to bind the CEP164 (Extended Data Fig. 3j, k), a high concentration of this protein can pass through the nuclear envelop (Extended Data Figs. 3l, m, q, r), translocate to the cilia base and induce a ciliopathy (Extended Data Figs. 3n-p), ultimately promoting the neurogenic responses of the Tis21-Luc reporter (Extended Data Fig. 3s).

### Microcephaly in a human post-mortem ZikV infected foetal brain is accompanied by a ciliopathy in ependymal cells

During development and upon termination of primary neurogenesis, monociliated NPCs differentiate into multiciliated ependymal cells (ECs)^42^. Hence, the ciliopathy described here in the chick embryo NPCs might persist in the ECs of ZikV infected human foetal brain. To address this hypothesis, we examined post-mortem tissue from a reported ZikV-infected microcephalic foetus from a ZikV infected mother. The pregnant woman reported possible exposure to the Aedes mosquito until the 12^th^ gestational week (GW) and ZikV RNA was detected in the GW22. After ultrasound and MRI imaging, the foetus was diagnosed with severe microcephaly and the pregnancy was terminated in the GW22+6. A severe developmental delay was evident in forebrain coronal sections, with almost complete cortical plate agyria when compared to the control sections of a non-pathological GW21 foetal brain. A thinner cortical plate (CP), and thinner intermediate (IZ) and subventricular zones (SVZ) were also detected (Figs. 4a, b). Infection of the tissue was confirmed by immunostaining for the viral NS5 protein^8^, which consistent with the viral infection early in gestation, spread from the ependymal layer, to the SVZ and CP (Fig. 4c). Moreover, the viral NS5 protein was detected in sub-ependymal cells (Fig. 4d) and in ECs, co-localizing with the acetylated α-tubulin-labelled cilia shaft (Figs. 4e, f), cilia that are much shorter than in the control GW 21 ECs (Fig. 4g).

**Figure 4.**
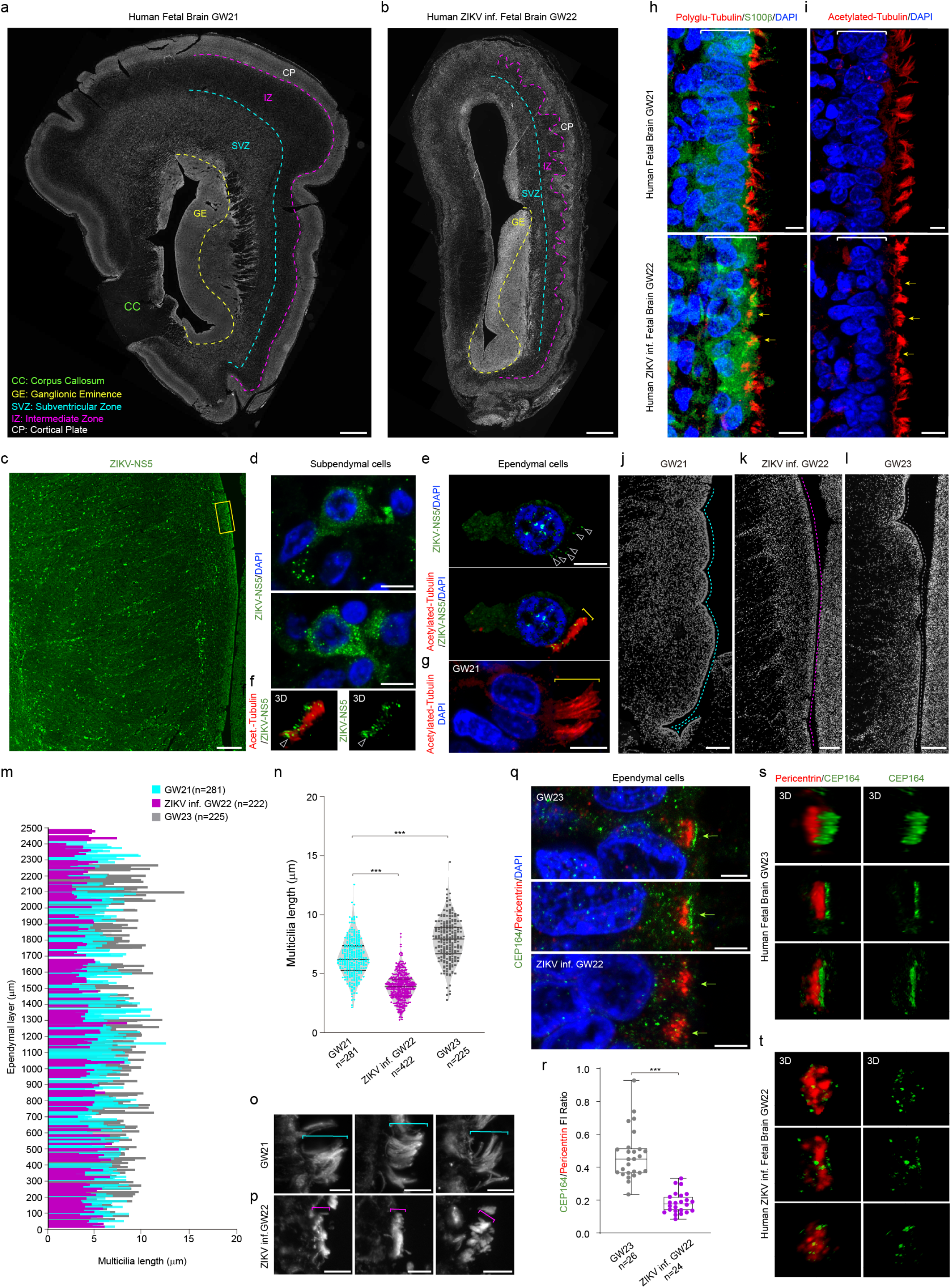
Microcephaly in ZikV infected Human foetal brain is accompanied by a ciliopathy in ependymal cells. **(a)** Representative image of control GW21 forebrain coronal sections indicating the different forebrain areas. **(b)** Representative image of ZikV infected GW22 forebrain coronal sections. **(c)** Representative immunostaining for the viral NS5 protein (green) in ZikV infectedGW22 coronal forebrain sections. **(d)** Selected images from c (yellow inset) showing the viral Zikv-NS5 (green) protein in subependymal cells with DAPI (blue) stained nuclei. **(e)** Selected images show acetylated-tubulin labelled multicilia (red) in ZikV-NS5 infected (green dots) ependymal cells with DAPI (blue) stained nuclei. Arrowheads point to the Zikv-NS5 protein and the bracket indicates the cilia length. **(f)** 3D image of ZikV-NS5 (green, arrowhead) localized to the basal acetylated-tubulin labelled multicilia. The arrowheads clearly show the green dots at the base of a cilia. **(g)** Selected image showing acetylated-tubulin labelled multicilia (red) in control GW21 ependymal cells with DAPI (blue) stained nuclei, in which the bracket indicates the cilia length. **(h)** Selected images of poly-glutamylated-tubulin stained multicilia (red) in S100β^+^ ependymal cells (green) of control GW21 foetal brain, and ZikV GW22 foetal brain. **(i)** Selected images of acetylated-tubulin stained multicilia (red) in control GW21 and ZikV GW22 ependymal cells in which the yellow arrows indicate multicilia shortening. **(j-l)** Selected images depicting the 2.5 mm ependymal areas (dotted lines) used to measure cilia length in control GW21, ZikV infected GW22 and control GW23 forebrain sections. **(m)** Plots of the cilia length along the ependymal layer at the developmental stages indicated. **(n)** Violin plots of the cilia length in ependymal cells of control GW21, ZikV infected GW22 and control GW23 forebrain sections (the upper and lower lines indicate the interquartile range, the middle line the median). **(o,p)** Selected images of acetylated-tubulin stained multicilia in control GW21 (**o**) and in ZikV infected GW22 ependymal cells (**p**, brackets indicate cilia length). **(q)** Selected images of pericentrin stained multiple centrosomes (red) and CEP164 (green) stained multicilia base in control GW21 and ZikV infected GW22 ependymal cells. **(r)** Box plot, median, quartiles (box) and range (whiskers) of CEP164/Pericentrin fluorescence intensity Ratio, showing reduced expression of CEP164 in the multiple centrosomes of ZikV infected ependymal cells. **(s,t)** 3D image showing Pericentrin (red) and CEP164 (red) localization to the base of multicilia in control GW23 (**s**) and ZikV infected GW22 (**t**) ependymal cells. \*\*\**p*<0.001, \*\**p*<0.01, \**P*<0.05. **n**, Kruskal-Wallis test, **r**, Mann-Whitney *U* test. Scale bars 1.5 mm (**a**, **b**), 200 μm (**c, j**-**l**). 6 μm (**d, e, g**-**i, o, p**), 3 μm (**q**).

In addition, we observed a severe disorganization of the S100β^+^ ECs lining the ventricle (Fig. 4h), as reported in ZikV infected macaque tissues^9^, with differentiating ECs expanding their apical surface and assembling multiple motile cilia^43^. To investigate whether ZikV infection affected ciliogenesis, the length of the acetylated α-tubulin cilia shafts protruding from the VZ surface was measured in the ~2.5 mm ependymal layer facing the Ganglionic Eminence (GE) in ZikV infected ECs, comparing these values to those in control ECs in the 21^st^ and 23^rd^ GW (Figs. 4i-l). While control tissue showed an increase in cilia length associated with EC maturation (GW21 median=6.3 ± 1.57 μm, n=281; GW23 median=7.8 ± 1.77 μm, n=225), the EC multicilia were significantly shorter and more disorganized in ZikV infected tissue (median=3.86 ± 1.14 μm, n=422: Figs.4m-p).

Multiciliated EC differentiation involves the assembly of a large number of centrioles that migrate apically and anchor to the plasma membrane. In the case of these cilia, Cep164 localizes to the distal appendages and Rootletin extends from the proximal end of the centrioles to their rootlets^43^ (Figs. 4q, s). To search for the mechanisms underlying the ZikV mediated ciliopathy in human tissue, we quantified the endogenous Cep164 protein in ECs, relative to the centriole marker Pericentrin. ZikV infected ECs had significantly less CEP164 protein than control ECs, which is itself required for motile cilia assembly^44^ (control mean=0.47 ± 0.15, n=26; ZikV infected mean=0.18± 0.06, n=24: Fig. 4 q, r). Indeed, 3D imaging highlighted the disturbed CEP164 distribution at the apical surface of ZikV infect ECs relative to the controls (Figs. 4s-t). Together our analysis revealed that ZikV infection of human foetal brain tissue not only compromises primary neurogenesis, resulting in a severe microcephalic brain but also, it disrupts EC differentiation and provokes a ciliopathy in these cells. This ciliopathy is likely to impair the flux of CSF elements necessary for brain homeostasis in these cells, as well as for toxin wash-out, further contributing to the developmental malformations associated with congenital Zika syndrome, such as microcephaly and ventriculomegaly.

## DISCUSSION

ZikV has evolved multiple mechanisms to exploit or perturb fundamental cellular processes in NPCs^17^. Previous data showed that NS4AB interferes with the mTOR pathway in NPCs, reducing proliferation and inducing autophagy^45^, and that NS2A interacts with AJs, while disrupting the integrity of the developing neuroepithelium and again, perturbing NPC proliferation^20^. Recently, the NS2B3 heterodimer was also shown to disturb the cell-cycle as it affects the host protein Septin-2 that is involved in NPCs cytokinesis^46^. To date, NS5 has only been shown to counteract host antiviral mechanisms by targeting the interferon pathway^47^. Here we describe a toxic effect of NS5 on NPCs by showing its capacity to interact with multiple host proteins at the cilium base, consequently promoting a ciliopathy and a premature NPC differentiation.

Primary and motile cilia differ in their structure, composition and function. In the developing brain, primary cilia are non-motile signaling organelles present on NPCs that help integrate growth signals. Multiple motile cilia are found on the surface of the ECs lining all the brain ventricles, where they contribute to the flow of CSF. During brain development, monociliated NPCs differentiate into multiciliated ECs, providing an ideal system to study the impact of pathogen infection on these two developmental stages (Fig. 5). The primary cilium has a microtubule-based core and an axoneme that extends from a specialized centriole at the base of the cilium, the site where Cep164, Rootletin and Bart are located (Fig. 1e). Differentiation of ECs involves the assembly of a large number of centrioles that migrate apically and anchor to the plasma membrane^43^ (Fig. 5). Here we show that in NPCs, ZikV-NS5 binds directly to the ciliary basal body, impairing the normal development of this organelle and resulting in cilia shortening (Fig. 1-2). It seems that NS5 hijacks Cep164, Rootletin and Bart and/or blocks the natural host protein-protein interactions in the cilia basal body, perturbing its natural assembly. Failure to elongate a primary cilium after mitosis diminishes the reponses to external stimuli like Shh, and to other growth factors that promote symmetric NPC division and CNS growth (Fig. 5)^33,34^. Remarkably, reduced levels of CEP164 and centriole damage were previously associated to ZikV infection^6^. In addition, dysregulation of neurogenesis through the Notch pathway, a modulator of NPCs response to Shh ligand, was also reported in ZikV infections^48^. Hence, the primary ciliopathy provoked by ZikV-NS5 seems to be responsible for the microcephaly induced by ZikV. Interestingly, we found that in huamn post-mortem tissue of a presumed ZikV-infected microcephalic fetus from a ZikV infected mother, ECs exhibited a severe ciliopathy that might prevent the flow of the CSF necessary to maintain brain homoeostasis and toxin washout (Figs. 4-5). The distinction between a motile ciliopathy and a primary ciliopathy seems to be unclear, as hydrocephalus may also be observed in a number of primary cilia disorders^49^. Indeed, here we show that a single pathogen can cause both types of ciliopathy, depending on the cell targeted.

**Figure 5.**
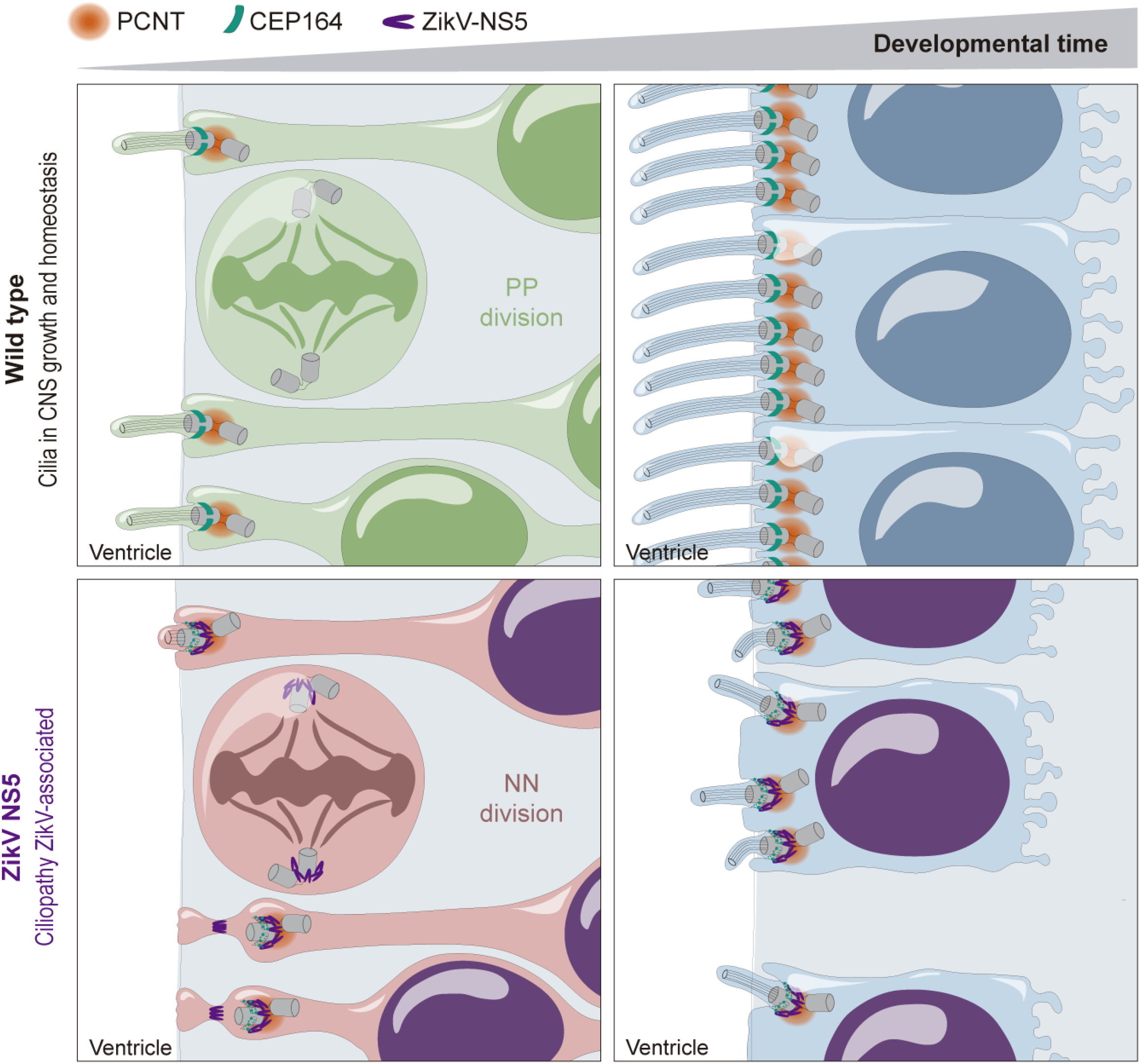
ZikV-NS5 protein causes ciliopathy. Monociliated NPCs divide symmetrically (PP divisions) to ensure normal growth in the ventricular zone (VZ). In NPCs, ZikV-NS5 interacts with host proteins at the base of the cilia, such as CEP164, to cause a ciliopathy and promote terminal neurogenic (NN) divisions. Upon termination of primary neurogenesis, NPCs differentiate into multiciliated ependymal cells (ECs) in which ZikV-NS5 conserves its interaction with the same host proteins at the base of the cilia to provoke a ciliopathy and the disruption of the ependymal layer (EL).

We have also demonstrated that both ciliopathy and the imbalance in the mode of division produced by ZikV-NS5 are strongly dependent of its quaternary structure. The recently reported X-ray structures of the full-length ZikV NS5 revealed a dimer based helicoidal arrangement of the protein that is mediated by MTase-MTase interactions, involving residues Y25, K28 and K29 (Fig. 3h)^13^. Here we show that the monomeric ZikV-NS5 Y25A/K28S/K28A variant is retained within the cell nucleus forming a diffuse pattern. This data suggests that nuclear multimerization of NS5 is required to escape from this cellular compartment and travel to the basal body. Indeed, our results show that the capacity of ZikV-NS5 to form multimers contributes to its interference with components of the cilium base, a fundamental mechanism underlying the distinct developmental malformations attributed to congenital Zika syndrome. These findings suggest alternative routes that could be followed in order to discover new antiviral agents that target the intermolecular interfaces involved in dimer/oligomer formation.

## Acknowledgements

The authors are indebted to Dr Elena Rebollo and Jaume Boix for their invaluable technical assistance at the AFMU Facility (IBMB); We thank Leica Microsystems for supporting and collaborating with the AFMU Facility (IBMB). We are grateful to researchers that kindly provided DNAs and antibodies, as indicated in the reporting summary. The work in EM’s laboratory was supported by grants BFU2016-77498-P, BFU2016-77498-P and La Maratò de TV3 foundation 201833-10. The work in NV’s laboratory was supported by grants BIO2017-83906-P, Maria de Maeztu Unit of Excellence MDM-2014-0435 (MCIU) and La Maratò de TV3 foundation 201833-10

## Author Contributions

**MS** conceived and performed most of the experiments, analysed the data, discussed the results and revised the manuscript.

**DSF** conceived and performed the structural and biochemical experiments, with the help of **VMRA**. analysed the data, discussed the results and revised the manuscript.

**JBA and EGG** contributed to the experiments, the image acquisition, image analysis and quantification, and the statistics, and they revised the manuscript.

**EMS and SRC** provided the human tissue.

**NV** conceived the structural experiments, discussed the results and revised the manuscript.

**EM** conceived the experiments, analysed the data, discussed the results and drafted the manuscript.

---

**Extended data Figure 1**

**(a)** Scheme showing the DNAs co-electroporated for immunohistochemistry.

**(b-e)** Subcellular localization of ZikV non-structural (NS) proteins co-electroporated with the cilia marker Arl13b-RFP (red) into stage HH12 chick embryos. At 16 hours post-electroporation (hpe) viral proteins were detected using an anti-FLAG antibody and visualized by confocal microscopy. In electroporated NPCs, ZikaV-NS1 (**b**) and ZikaV-NS2 (**c**) are located in the cytoplasm, excluded from the nucleus, the FOP labelled (purple) centrosome and the Arl13b-RFP (red) labelled cilia. ZikV-NS3 (**d**) has a cytoplasmic distribution and it accumulates at the cilia base. ZikV-NS4 (**e**) localizes at the membrane and it is excluded from the FOP labelled centrosome.

(**f**) Scheme showing the DNAs co-electroporated for immunohistochemistry. Representative images of spinal cord sections from chick embryo (stage HH12+24hpe), showing the electroporation of control (red) of ZikV-NS5 (red), Tubb3enh (green) and HUC/D (blue) immunostaining of differentiated neurons.

**(g)** Representative images of spinal cord sections from a chick embryo (stage HH12+24hpe), showing ZikV-NS5-FLAG (green) electroporation and anti-PH3 immunostaining (red). **(h)** Quantification of pH3^+^ cells in the electroporated control vs ZIKV-NS5 electroporated NT (Ctr median=10.5 ± 1.3 vs ZIKV-NS5 median= 10 ± 0.9 pH3^+^ cells, n=5 embryos)

**(i)** Representative images of spinal cord sections from a chick embryo (stage HH12+24hpe), showing ZikV-NS5-FLAG (green) electroporation and anti-Caspase 3 immunostaining (red). **(j)** Quantification of Caspase 3^+^ cells in the electroporated control and ZIKV-NS5 electroporated NT (Ctr median=3 ± 2.2 vs ZIKV-NS5 median=3.5 ± 1 Caspase 3^+^ cells, n=5 embryos)

**(k-n)** ZikV-NS5 localizes to the centrosomes at different mitotic phases.(**k**) ZikV-NS5-FLAG revealed by anti-FLAG staining (green) at centrosomes labelled with anti-FOP (purple) at prophase. DAPI (blue) labels the chromosomes and the arrows point to the centrosomes. **(l)** ZikV-NS5-FLAG (green) localizes symmetrically to the centrosomes during metaphase. **(m,n)** ZikaV-NS5-FLAG (green) localizes to centrosomes during anaphase and telophase. The mother centrosome is labelled with anti-polyglutamylated Tubulin (red) and Arl13b (red).: Scale bars 5 μm).

**(o-q)** Scheme showing the DNAs co-electroporated for immunohistochemistry (**o**). Subcellular localization of ZikV NS1-4 proteins co-electroporated with ZikV-NS5-GFP (green) and with the cilia marker Arl13b-RFP (red) in stage HH12 chick embryos. At 16 hpe viral proteins were detected using anti-FLAG antibody and visualised by confocal microscopy. In electroporated NPCs, ZikV-NS5 (green) remained associated to the cilia base identified by Arl13b-RFP (red)

**(r)** Sequence alignment highlighting the conservation between human and chicken proteins in the ZikaV-NS5-interaction domain (red box)

ns: not significant. **h**-**j**, Mann-Whitney *U* test. Scale bars 9 μm (**b**-**e**), 2 μm (inset **b**-**e**, **q**), 75 μm (**f**), 70 μm (**g**, **i**), 5 μm (**k**-**n**), 6 μm (**p**)

**Extended data Figure 2**

**(a)** The scheme depicts the cilia length in relation to cell cycle phases of dividing NPCs, with the microtubules in cilia highlighted in red.

**(b)** Selected images show Polyglutamylated-tubulin labelled cilia (red) and FOP stained centrosomes lining the NT lumen (dotted line), in control and ZikV-NS5-FLAG electroporated embryos (green, arrow pointed)

**(c)** Violin plots of the cilia length in control and ZikV-NS5 electroporated NPCs (the upper and lower lines indicate the interquartile range, the middle line the median)

**(d,e)** Selected images show ZikV-NS5-FLAG (purple arrow) co-localized to FOP stained centrosomes (red) and to the CEP164 stained mother centriole (green arrow, **d**) or Rootletin staining (green arrow, **e)**

**(f)** *En face* images of N-cadherin labelled AJs (purple), endogenous CEP164 (green) labelled centrosomes and ZikV-NS5-FLAG subcellular localization (red arrow)

**(g)** *En face* images of N-cadherin labelled AJs (purple), endogenous Rootletin (green) labelled centrosomes, and ZikV-NS5-FLAG subcellular localization (red arrow). The yellow arrow points to a centrosome where CEP164 and Rootletin expression is reduced relative to the neighbouring NPCs.

**(h)** *En face* images of endogenous apical BART (green) and ZikV-NS5-FLAG subcellular localization (red). Green lines and yellow arrows point to BART and ZikV-NS5 de-localization from the centrosome towards the apical belt, respectively. Centrosomal ZIKV-NS5 is highlighted with a yellow dashed circle.

**(i)** *En face* images of N-cadherin (purple) and FOP stained centrosomes (red) in wild-type embryos. Lateral positioned centrosomes are correlated to the apical area restriction.

**(j)** Representative images of ZikV-NS5-FLAG (purple) co-localization with Arl13b (red) and with endogenous BART (green) lining the NT lumen. **(k)** Boxed areas in j, green arrows point to BART localization at the apical belt, red arrows to Arl13bshort cilia, purple arrows to ZikV-NS5 at the centrosome and apical abscising area.

**(l)** Violin plots of the ratio of the apical area between control and ZikV-NS5 transfected cells, and surrounding non-transfected neighboring cells (the upper and lower lines indicate the interquartile range, and the middle line the median).

**(m,n)** *En face* and 3D images of a NPC where N-cadherin labelled AJs (purple), FOP labelled centrosome (red) and the ZikV-NS5-FLAG subcellular localization (green). The purple arrow points to the apical area, the red arrow to a laterally positioned centrosome, the green arrow to ZikV-NS5 at the centrosome.

**(o)** Representative images of ZikV-NS5-FLAG (purple) electroporated NPCs in which ZikV-NS5-FLAG was located at the cilium base, as determined by Arl13b-GFP (green arrows) and correlated with pTis21-RFP (red) expression.

**(p)** Scheme of electroporated DNAs and quantification of the proportion of pTis21-RFP^+^ cells, according to the subcellular distribution of ZikV-NS5 (the error bars correspond to the mean ± s.e.m.)

\*\*\**p*<0.001, \*\**p*<0.01; exact *P* values 0.0011 (**l**), *\*P*<0.05, ns: not significant. **c**, **l** Mann-Whitney *U* test; **p**, one-way ANOVA. Scale bars 2 μm (**b**, **d**, **e**, **k**), 4 μm (**f**-**i, m**). 7 μm (**j, o**).

**Extended data Figure 3**

**(a)** The scheme depicts the MTase (light blue) and RdRp (dark blue) domains in the ZikV-NS5 protein, with the linker highlighted in yellow. 3D representation of one ZikV-NS5 monomer showing an orientation from the top of the RdRP, with the MTase domain in grey, and the RdRP fingers, palm and thumb domains in blue. Red indicates the position of the NLS.

**(b)** Quantification of the Luc/Renilla activity of the pTis21-Luc reporter after electroporation of the DNAs indicated (the plots show the mean ± s.e.m., n=6-8 embryos/condition)

**(c)** Scheme representing the reporter co-electroporation experiments, harvested at 24 hpe for either flow cytometry analysis or for anti-PH3 immunostaining.

**(d)** Selected images of pSox2^+^ (green) and pTirs21^+^ (red) NPCs, and pH3 stained mitoses (blue) in the control or ZikV-NS5 NT.

**(e)** Quantification of reporter-expressing pH3^+^ dividing cells in each condition: PP (pSox2^+^/pTis21^−^, green), PN (pSox2^+^/pTis21^+^, yellow) and NN (pSox2^−^/pTis21^+^, red). The data represents the mean ± s.e.m)

**(f)** Quantification of reporter-expressing FACS sorted cells in each condition, the data representing the mean ± s.e.m.)

**(g)** Representatives *en face* images of ZikV-NS5-FLAG ring-like aggregates (green). The nucleus is highlighted with a dashed-dotted line

**(h,i)** Representatives *en face* images of ZikV-NS5-FLAG (green) where the yellow arrow follows the basal to apical ring-like distribution of the protein all along the apical foot **(h)** and at the apical-end foot at the centrosome level (**i**). N-cadherin labelled AJs (purple) and the FOP labelled centrosome (red).

(**j,k**) HEK-293 cells were co-transfected with WT or MultimDead ZikV-NS5 and the DNAs indicated, and the cell extracts were analysed by western blots (**j**). Quantification of Pull-down (PD) assays (**k**) shows a less efficient binding of the MultimDead ZikV-NS5 to Cep164 and Emerin.

**(l-o)** Representative images of the MultimDead ZikV-NS5 subcellular localization associated with Arl13b labelled cilia (red) and FOP labelled centrosomes (blue). Ciliopathy (red arrow) associated with the MultimDead ZikV-NS5localization to the cilium base (green arrow). MultimDead ZIKV-NS5 was also detected in the tip of cilia (yellow arrow).

**(p)** Violin plots of the cilia length in association with the subcellular localization of the MultimDead ZikV-NS5(the upper and lower lines indicate the interquartile range, the middle line the median).

**(q,r)** Representative images of ZikV-NS5-FLAG nuclear aggregates (green arrow) associated with increasing electroporation. **(r)** MultimDead ZikV-NS5 only causes nuclear envelope disruption at high concentrations (green arrow).

**(s)** Quantification of the Luc/Renilla activity of the pTis21-Luc reporter after electroporation of the DNAs indicated (the plots show the mean ± s.e.m., n=6-8 embryos/condition).

\*\*\**p*<0.001, \*\**p*<0.01; exact *P* values 0.0015 (**e**) 0.0044 (**f**) 0.0051 (**s**), *\*P*<0.05; exact *P* values 0.049 (**e**), ns: not significant. **b**, **e**, **f, s** one-way ANOVA, **p**, Kruskal-Wallis test. Scale bars 30 μm (**d**), 3 μm (**g-i, n, o**). 5 μm (**l, m, q, r**).

**Supplementary Figure 1**

Source data of Western Blots with size marker indications in KDa. Dashed line boxes mark the cropped area.

**Supplementary Figure 2**

Source data for Flow Cytometry Report Summary

**Supplementary Table 1**

ZikV-NS5-host interactorme determined by Yeast two hybrid and subdivided based on their subcellular localization according to the Gene Ontology (GO) analysis

**Supplementary Table 2**

Comparative analysis of the ZikV-NS5 yeast two hybrid data with two independant interactomes previously published

**Supplementary Table 3**

List of Antibodies used in this study

**Supplementary Table 4**

List of oligonucleotides used in this study

**Video 1**

3D reconstruction of a ZIKV-NS5-Flag (shown in Green) electroporated NPC showing a premature basal migration of the Fop+ centrosome (shown in Red). The Apical area of the electroporated NPC is delimitated with N-Cadherin (shown in Magenta)

**Video 2**

3D reconstruction of a ZIKV-NS5-Flag (shown in Green) electroporated NPC showing a short apical foot with a restricted apical-end foot area (Yellow Arrow) delimitated with N-Cadherin (shown in Magenta, asterisk points the nucleus)

**Video 3**

3D reconstruction of a ZIKV-NS5-Flag (shown in Green) electroporated twos neighbors NPCs showing a short apical foot with a restricted apical-end foot area (Yellow and blue Arrows) delimitated with N-Cadherin (shown in Magenta, Asterisks with the same color code than the arrows point the corresponding nucleus)

**Video 4**

3D reconstruction of a ZIKV-NS5-Flag (shown in Green) electroporated NPC showing a short apical foot with a restricted apical-end foot area (Yellow Arrow) delimitated with N-Cadherin (shown in Magenta, asterisk points the nucleus)

**Video 5**

3D reconstruction of a Dsredex (shown in Green) electroporatde NPC showing a long apical foot with a restricted apical-end foot area (Yellow arrow) delimitated with N-Cadherin (shown in Magenta, asterisk points the nucleus).

**Video 6**

3D reconstruction of a Dsredex (shown in Green) electroporated NPC showing a long apical foot with a restricted apical-end foot area (Yellow Arrow) delimitated with N-Cadherin (shown in Magenta, asterisk points the nucleus).

