## Supplementary figures and images for "Multimerization of Zika Virus-NS5 causes a ciliopathy and forces premature neurogenesis"

### Extended Figure 1

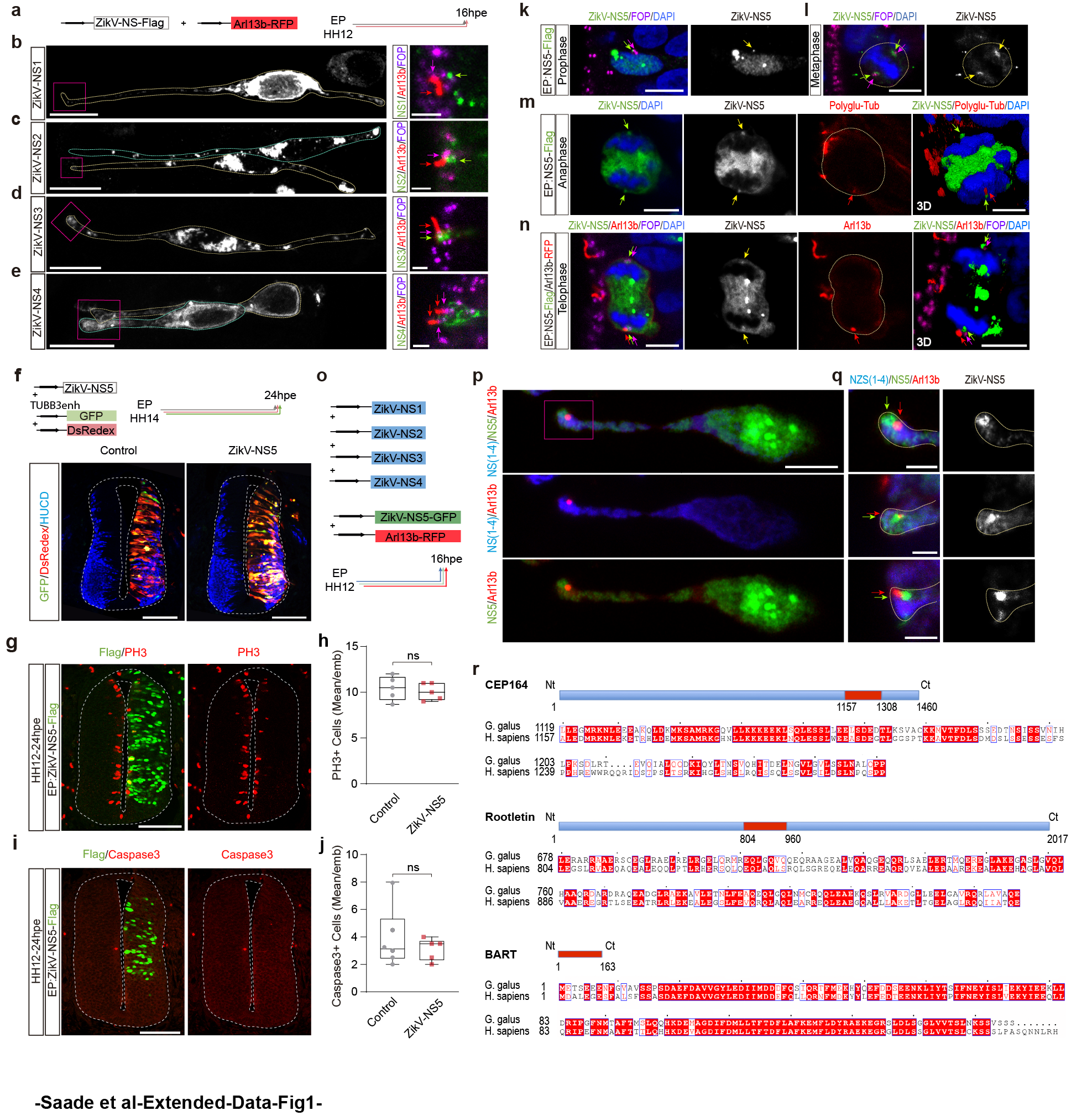

### Extended Figure 2

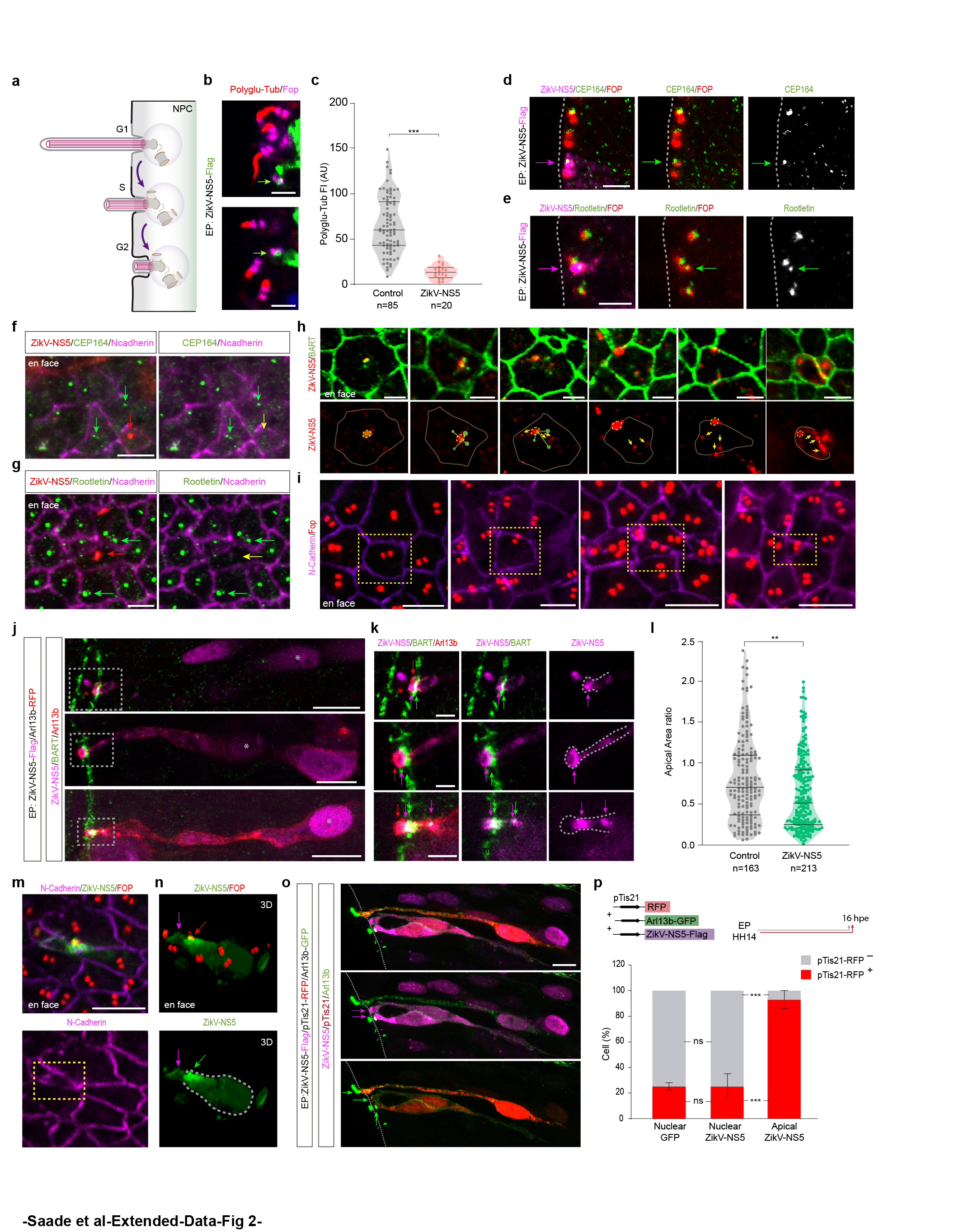

### Extended Figure 3

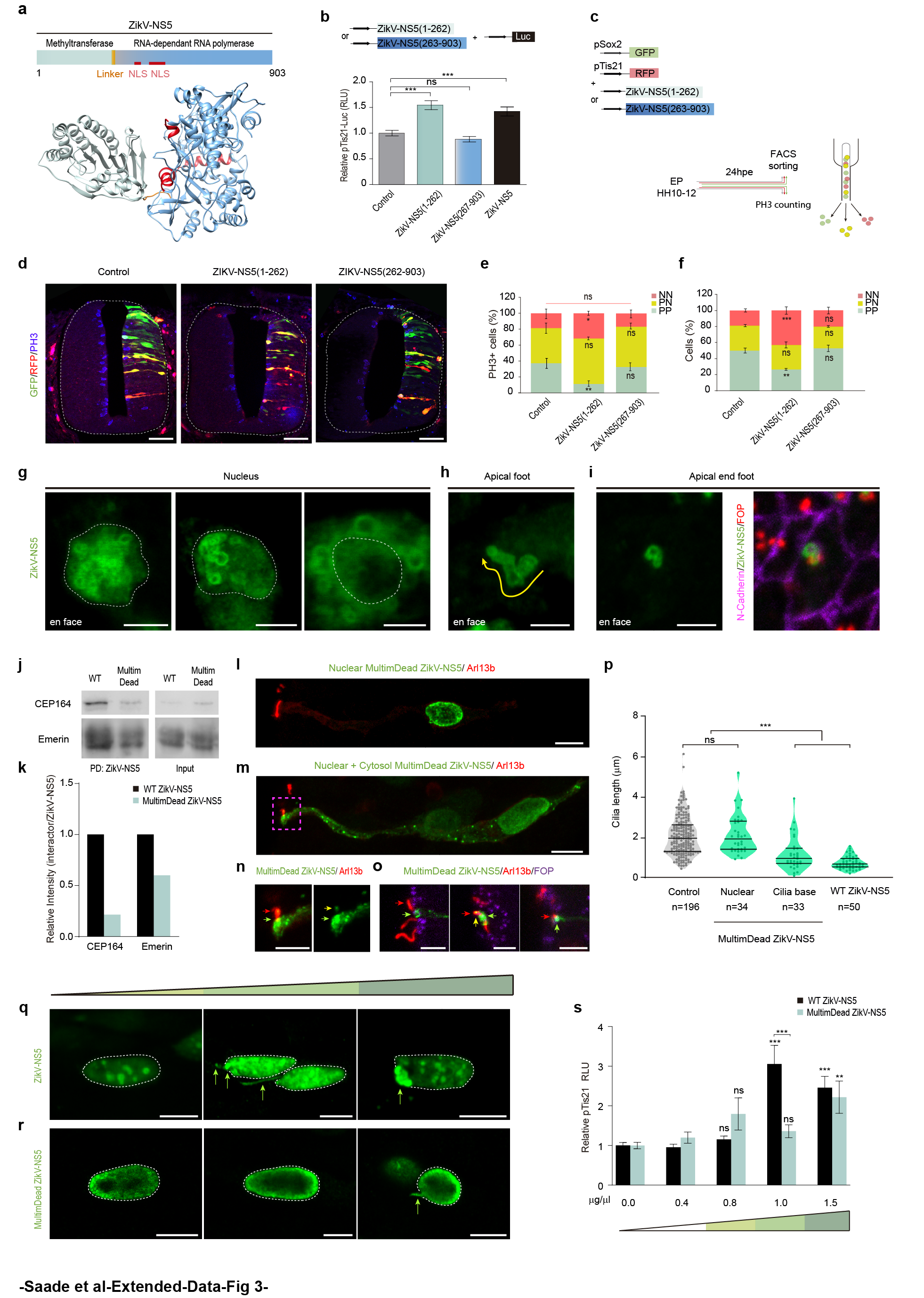
